# Cell binding, uptake and infection of influenza A virus using recombinant antibody-based receptors

**DOI:** 10.1101/2024.07.29.605726

**Authors:** Oluwafemi F. Adu, Milagros Sempere Borau, Simon Früh, Umut Karakus, Wendy S. Weichert, Brian R. Wasik, Silke Stertz, Colin R. Parrish

## Abstract

Human and avian influenza A viruses bind to sialic acid (Sia) receptors on cells as their primary receptors, and this results in endocytic uptake of the virus. While the role of Sia on glycoproteins and/or glycolipids for virus entry is crucial, the roles of the carrier proteins are still not well understood. Furthermore, it is still unclear how receptor binding leads to infection, including whether the receptor plays a structural or other roles beyond being a simple tether. To enable the investigation of the receptor binding and cell entry processes in a more controlled manner, we have designed a protein receptor for pandemic H1 influenza A viruses. The engineered receptor possesses the binding domains of an anti-HA antibody prepared as a single chain variable fragment (scFv) fused with the stalk, transmembrane and cytoplasmic sequences of the feline transferrin receptor type-1 (fTfR). When expressed in cells that lack efficient display of Sia due to a knockout of the *Slc35A1*gene which encodes for the Solute Carrier Family 35 transporter (SLC35A1), the anti-H1 receptor was displayed on the cell surface, bound virus or hemagglutinin proteins, and the virus was efficiently endocytosed into the cells. Infection occurred at similar levels to those seen after Sia reconstitution, and treatment with clathrin-mediated endocytosis (CME) inhibitors significantly reduced viral entry.

**IMPORTANCE.:** Influenza A viruses mostly circulate among avian reservoirs, and also can jump hosts to cause epidemics in mammals, including among humans. A key interaction of the viruses is with host cell Sia, which vary in chemical form, in their linkages within the oligosaccharide, and in the attachment to surface glycoproteins or glycolipids with different properties. Here we report a new method for examining the processes of receptor binding and uptake into cells during influenza A virus infection, by use of an engineered HA-binding membrane glycoprotein, where an antibody is used as the binding domain and the transferrin receptor uptake structures mediate efficient entry, which should allow us to test and manipulate the processes of cell binding, entry, and infection.

## INTRODUCTION

Influenza A viruses (IAVs) are negative sense single-stranded segmented RNA viruses of the Orthomyxoviridae family. They have an envelope that anchors two important glycoproteins, hemagglutinin (HA) and neuraminidase (NA), which mediate infection and tropism of the virus (1–3). For most IAVs, HA facilitates entry into competent cells by binding to sialic acids (Sia), often in the form of N-acetylneuraminic acid (Neu5Ac) as the terminal glycan of an oligosaccharide linked to either surface glycoproteins or glycolipids (4–6). Bat IAVs (H17 and H18) and avian H19 are an exception to Sia-mediated entry, as those use MHC class II glycoproteins for attachment and uptake (1, 7–9).

Host-specific evolution of the IAV HA leads to observed selective binding of differently linked Sias. Human IAVs favor Sia with an α2,6 linkage to a neighboring galactose or GalNac residue, while avian IAV strains favor α2,3 linked Sia (10–12). Additionally, the neuraminidase (NA) plays a significant role in facilitating IAV infection in animals due to its ability to cleave Sia, thereby reducing virus clustering during viral budding and release. NA also modifies non-productive HA-Sia interactions with mucus within the respiratory tract (2). This, in turn, promotes productive viral attachment and infection of the underlying epithelial cells. Furthermore, NA has been shown to sometimes also mediate Sia binding, and to also directly interact with the immunomodulatory cell adhesion molecule CEACAM6 (CD66c) to promote IAV entry into competent cells (13). However, whether this CD66c interaction is independent of Sia is unknown. In total, for most virus:host combinations there is holistic HA-NA-Sia interaction balance that is necessary to promote IAV fitness (2).

Following Sia binding, viral entry involves significant levels of clathrin-mediated endocytosis (CME) (14–16). However, it is still unclear how Sia binding efficiently leads to entry through that pathway, and also whether it involves the preferential use of Sia expressed on specific surface glycoproteins or glycolipids (4, 17, 18). It is theorized that attachment to Sia alone may not be sufficient to trigger signaling cascades necessary for effective internalization, and that efficient endocytosis involves receptor crosslinking by the multivalent influenza virus particle, which also increases the avidity of binding (19).

Multiple Sia-dependent receptors or co-receptors have been suggested for IAVs. These include nucleolin (20, 21), the calcium voltage pump CaV1.2 (22–24) and NK cell specific NKp44/46(25, 26). These receptors have been suggested to aid in virus attachment by directly interacting with the HA. Other receptor candidates include members of the receptor tyrosine kinases such as the epithelial growth factor receptor (EGFR) (19) or the G-protein coupled receptors (GPCRs) like, β-arrestin, and the free fatty acid receptor 2 (FFAR2) (27, 28). Those are thought to engage (likely indirectly) with the virus upon uptake. Other alternative pathways for IAV entry may involve the Sia present on N-linked-glycans on the viral glycoproteins binding to different C-type lectins expressed on various immune cells including dendritic cells (DC-SIGN) (29), macrophages (MMR and MGL) (30) or by Langerhan cells in the human airway (Langerin)(31). Those interactions may play direct or indirect roles in IAV uptake and internalization leading to infection.

Understanding the specific interactions between IAV and many of the above-described candidate receptor components and their roles in directing viral entry and infection has been challenging. Several factors contribute to this difficulty, including the presence of different IAV strains that may engage distinct co-receptors, diverse tropisms for various cell types which may involve different host cell receptors or co-receptors, possible redundancy in the entry mechanisms of the virus, and difficulties associated with analyzing and altering the glycan-protein expression in a predictable way. In addition, methods currently employed may lack specificity and sensitivity, and *in vitro* systems may not fully recapitulate the *in vivo* processes in the respiratory or gastrointestinal tracts of the natural hosts.

To provide new controllable methods for studying IAV entry and uptake mechanisms, we have developed a novel antibody-based receptor (ABR) chimera that mediates pandemic H1N1 entry/infection without the involvement of Sia. This allows for more flexibility and manipulation of the binding, entry and infection processes in different systems and cell types. We show that a chimeric glycoprotein receptor rescued pandemic H1N1 entry and infection in both HEK and A549 cells that lacked cell surface Sia expression.

## MATERIALS AND METHODS

### Cells and viruses

HEK 293 (HEK) and A549 cells were obtained from the American Type Culture Collection (ATTC) and were cultured at 37°C, 5% CO_2_. HEK *Slc35A1* knock out (KO) cells were prepared as previously described in (5). A549 *Slc35A1* KO cells were generated by CRISPR-Cas9-mediated genome editing. Briefly, A549 cells were reverse-transfected with pre-assembled ribonucleoproteins (RNP) consisting of Alt-R S.p. Cas9 nuclease (IDT) in complex with Alt-R CRISPR-Cas9 crRNA that targets exon 1 of the *Slc35A1* gene (*Slc35A1*_crRNA: (AltR1) rGrA rCrCrC rArGrU rUrCrU rCrArC rCrUrC rUrCrG rGrUrU rUrUrA rGrArG rCrUrA rUrGrC rU (AltR2) (IDT) and Alt-R CRISPR-Cas9 tracrRNA – ATTOTM 550 (IDT), using RNAiMax (Thermo Fisher Scientific). At 48h post transfection, single cell clones were generated by limiting dilution in 96-well plates and screened by next generation sequencing (NGS). Genomic DNA was extracted from single cell clones using QuickExtract DNA Extraction Solution 1.0 (Lucigen) and the region of interest was amplified by two consecutive PCRs. First PCR was run with primers containing adapters for TruSeq HT index primers (indicated by underlined nucleotides): SLC35A1_frw: 5′-CTT TCC CTA CAC GAC GCT CTT CCG ATC TTC TAT GAC CAC AAG GGG CGG TC-3’ and SLC35A1_rev: 5’-GAC TGG AGT TCA GAC GTG TGC TCT TCC GAT CTA GCG GCT CCA CGC AAA CTC C-3’. The second PCR was run using TruSeq HT index primers D50x and D70y to generate barcoded amplicons. After gel extraction, barcoded amplicons were analyzed by MiSeq (Illumina). A genotypically *Slc35A1* KO clone was selected and phenotypically validated by Sambucus nigra lectin (SNA) and Maackia Amurensus II staining (see below for method). All cells used in this study were grown and maintained in Dulbecco’s modified Eagle medium (DMEM, Gibco) with 10% fetal calf serum (FCS) and 50μg/ml gentamicin (HEK) or 100 U/mL Penicillin, and 100μg /mL Streptomycin (A549) (Gibco).

A/California/04/2009 (Ca’09) was derived from reverse genetics plasmids as previously described (32). A/Netherland/602/2009 and Neth/09-*Renilla* were generated following previously described protocols (33)

### Lectin staining

Flow Cytometry analysis: HEK cells were grown to sub-confluency in a 12-well dish, collected using Accutase (Sigma), and blocked with Carbo-Free Blocking Solution (Vector Laboratories). Cells were incubated with FITC-conjugated SNA/MAL I (Vector Laboratories) for 30 minutes on ice, and signal intensities for 10,000 cells were assessed using a Millipore Guava EasyCyte Plus flow cytometer (EMD Millipore). Data was analyzed with FlowJo software (TreeStar).

For A549 cells, the cells were washed with PBS, detached with 0.25% trypsin-EDTA (ThermoFisher Scientific), resuspended in FACS buffer (PBS with 2% BSA and 1mM EDTA), and incubated with biotinylated SNA or MAL II lectins (Vector Laboratories) for 1 hour at 4°C. After washing, cells were stained with APC-Streptavidin (Biolegend) and LIVE/DEAD™ Fixable Near-IR Dead Cell Stain (ThermoFisher Scientific) for 30 minutes at 4°C. APC signal intensities of at least 5000 live cells were analyzed using a BD FACSVerse flow cytometer with BD FACSuite software. Data was analyzed in FlowJo. For both cell types, background signal intensities were subtracted, and values were normalized to their wild-type versions using GraphPad Prism.

Immunofluorescence Analysis: HEK cells were seeded into poly-D-lysine-treated glass slides (Invitrogen), fixed with 4% paraformaldehyde, blocked with Carbo-Free Blocking Solution (Vector Laboratories) and stained with FITC-conjugated SNA/MAL I lectins. Images were acquired using a Zeiss AxioSkop HBO 50 fluorescence microscope.

For A549 cells, 120,000 cells per well were seeded onto glass coverslips and incubated for 24 hours at 37°C. Cells were washed with PBS, fixed with 4% PFA, blocked with PBS containing 2% BSA, and stained with biotinylated SNA or MAL II lectins. Samples were then stained with Alexa Fluor 647-Streptavidin (Invitrogen) and DAPI (Sigma-Aldrich), mounted using ProLong Gold Antifade Mountant (Thermo Fisher Scientific), and imaged using a Leica SP8 confocal laser scanning microscope with a 63x objective. Images were acquired with LAS X software and processed with LAS AF Lite.

### Construction of chimeric receptors and HA-hFc probe

Structures of 5J8 Fab complexed with IAV-HA and details of their interactions were obtained from Protein data bank (PDB: 4M5Z)(34). The chimeric antibody-based receptor (fTfR-5J8) was prepared from the fTfR cytoplasmic, transmembrane and stalk domains (residues 1-122) (35) linked to the human monoclonal antibody (MAb) 5J8 (34, 36) single-chain variable fragments (scFv). The fTfR-5J8 receptor was prepared by in-frame cloning of the heavy and light chain variable sequences linked by a poly-GS-linker sequences (Fig. 1A-E) and expression from the pcDNA3.1 (-) vector under the control of the CMV immediate early promoter (37). The intact fTfR in the same expression vector was also used as control (35, 38).

**Figure 1.**
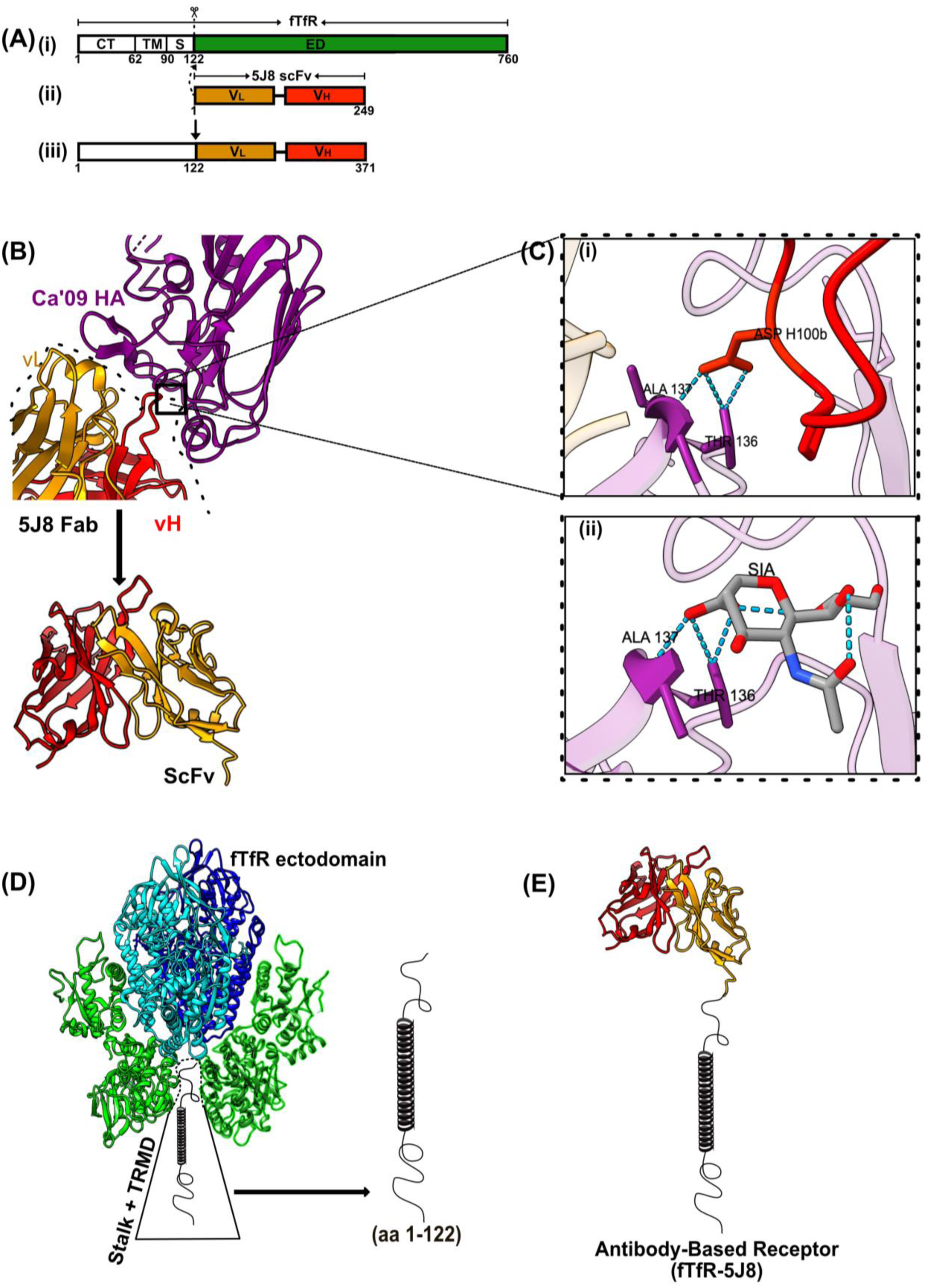
Design of Antibody-Based IAV receptor. (A) Cartoon showing the construction and cloning of the fTfR-5J8 chimera. (i-ii) Heavy and Light variable chains of MAb 5J8 were cloned as scFv constructs in-frame onto residues 1-122 of the feline transferrin receptor (fTfR) (iii) Receptor construct. (B) Crystal structure of Fab 5J8: A/Ca/07/2009-HA1 (PDB: 4M5Z). Purple (HA), Gold (Fab vL) and Red Fab vH) (B) Magnified H-bond interactions of HCDR3 (red) Asp 100 interactions with (i) Ala 137 and Thr136 HA residues (ii) Identical H-bond interactions of Sia with Ala 137 and Thr136 HA residues (PDB:3UBE) (34) (C) Cartoon of the TfR structure showing the ectodomain with Transmembrane (TRMD) and stalk domains (aa 1-122). (D) Cartoon of antibody based-receptor monomer (scFv from 5J8 Fab fused to the stalk and transmembrane and cytoplasmic domains of the fTfR). GS: Glycine-Serine Linker. Colors in the fTfR are similar to the different domains as defined for the human TfR; green (bottom half), protease-like domain; green (top half), apical domain, cyan and blue, helical domain (35).

To produce a probe for HA binding, the H1N1 Ca’09 HA ectodomain was fused via its C terminus in frame to the human IgG1 Fc, and the Gp64 baculovirus secretion peptide fused to the N-terminus as described previously (39, 40). Genes were synthesized and cloned into pFASTBAc-1 (Life technologies) to generate recombinant bacmids according to the manufacturer’s protocol, and as described previously (40).

Recovered recombinant baculoviruses obtained from Bacmid transfected Sf9 insect cells were used to infect suspension High Five^TM^ cells (Invitrogen). Two days post infection the proteins were purified by binding to HITrap ProteinG HP 1-ml column (Cytiva) and eluted with 0.1M glycine-HCL (immediately neutralized by 1M Tris, pH 9.0) by using an ÄKTA fast protein liquid chromatography (FPLC) system (GE Healthcare Life Sciences). Eluted fractions were dialyzed in PBS and concentrated using a 30-KDa Amicon filter tubes (EMD Millipore). Protein concentration was determined by Qubit (Invitrogen) and stored at -80°C in aliquots.

pMslc expressing the *human Slc35A1* gene was a gift from Dr. Christopher Buck (Addgene plasmid #32095). Plasmid stock was transformed into competent DH10B cells, blasticidin selected and plasmid DNA extracted using the E.Z.N.A. Plasmid DNA Mini Kit I (Omega BIO-TEK).

### Receptor expression

Chimeric receptors were transiently expressed in HEK *Slc35A1* KO cells by transfection of plasmids with Transit-X2 dynamic delivery system (Mirus Bio) 24hrs prior to virus (Ca’09) infection or HA-Probe binding assay. A549 *Slc35A1 KO* cells stably expressing the receptor constructs were generated by transduction with pLVX-fTfR-IRES-puro, pLVX-fTfR-5J8-IRES-puro or pLVX-Slc35A1-IRES-Neo-encoding lentiviruses in the presence of 8µg/mL polybrene. At 48 hours post-transduction, cells were subjected to selection with 1µg/mL puromycin or 1mg/mL neomycin.

### Analysis of receptor expression via qPCR

RNA was extracted from the indicated cell lines using the ReliaPrep RNA Miniprep kit (Promega) and then reversed transcribed into cDNA using oligo (DT) primers and Superscript IV RT (Promega) as per manufacturer’s instructions. RT-qPCR was performed with PowerTrack™ SYBR™ Green (Thermo Fisher Scientific) on a 7300 real-time PCR system (Applied Biosystems). The following primers targeting 5J8 and the extracellular domain of fTfR were utilized: fTfR fwd TGGCTGTATTCTGCTCGTGG, fTfR rev GCACTGATGTTTTCCTGGCG, 5J8 fwd GAAGGGGCTGGAGTGGATTG, 5J8 rev ATGCAGTATCGGGGTAACCA. GAPDH was included as an internal control. Samples where no amplification was detected were assigned a Ct of 40.

### Binding analysis with HA-Probe and A/Netherand/602/09 IAV

HEK WT, fTfR-5J8 or full-length fTfR expressing HEK *Slc35A1* KO *cells* were fixed, blocked with Carbo-Free blocking solution for 30mins (Vector) and incubated with the HA-hFc probe for 1hr at RT diluted in blocking solution. Bound HA-hFc was detected with anti-human IgG Fc Alexa 488 (Jackson Labs) and quantified using a Millipore Guava EasyCyte Plus flow cytometer (EMD Millipore), and data analyzed using the FlowJo software (TreeStar). Microscopic data was obtained by identical staining of receptor expressing or HEK WT cells in a 12 well plate (pre-treated with poly D Lysine (Invitrogen)) and images were taken using a Zeiss AxioSkop HBO 50 florescence microscope.

To assess IAV binding to the fTfR-5J8, A549 cells stably expressing *Slc35A1*, 5J8-fTfR or fTfR alone were inoculated with A/Netherlands/602/09 (A/Neth/09) at an MOI 25 for 1.5h on ice. Following two washes with PBS, cells were stained with LIVE/DEAD™ Fixable Near-IR Dead Cell Stain (Thermo Fisher Scientific) as per manufacturer’s instructions. Thereafter, cells were fixed and incubated with anti-HA 30D1 antibody (41) in FACS buffer. Bound virions in live cells were detected with anti-m-Alexa Fluor 647 (Thermo Fisher Scientific) and the signal intensities quantified using a Becton Dickinson FACSVerse flow cytometer. Flow cytometry data was processed with FlowJo v10.8.

### Cell infection and quantification

A/California/04/2009: HEK WT or HEK *Slc35A1* KO cells transiently expressing fTfR-5J8 or fTfR or the SLC35A1 were washed and incubated with the virus at an MOI of 0.2 for 1hr at 37°C. Growth media was added to each well and infection was allowed to proceed for another 7hrs. Cells were fixed with 4% PFA for 10 min, and then incubated with mouse anti-nucleoprotein (NP) antibody in permeabilization buffer (made in house), washed with PBS and further stained with anti-mouse Ig Alexa 488 for 1hr to detect viruses. DAPI stain was used to visualize cell nuclei. Imaging was done via the Zeiss AxioSkop HBO 50 florescent microscope, and images analyzed to obtain relative infected cells counts by ImageJ (NIH) (42).

### Neth/09-*Renilla* virus

A549 *Slc35A1* KO cells stably expressing SLC35A1 or the receptors were inoculated with Neth/09-*Renilla* virus as in (32). Cells seeded in 96 well plates were washed once with PBS and inoculated with Neth/09-*Renilla* virus at an MOI of 3 for 1 hour at 37°C. Following inoculum removal, cells were washed once with PBS and maintained in DMEM supplemented with 0.1% FBS, 0.3% BSA, 20 nM HEPES, 1% P/S (post-infection DMEM) and 6µM Renilla luciferase substrate (EnduRen, Promega). The mean relative light units (RLUs) for each time point were plotted and the area under the curve analyzed with GraphPad Prism 9.3.

### Endocytic entry

To assess the route of IAV entry, A549 cells, LV-S*lc35A1* or LV-fTfR-5J8 were pre-treated with Dynasore (40-20µM, Selleckchem), Pitstop 2 (15-10µM, Sigma-Aldrich), NH_4_Cl (25mM) or DMSO (0.15%) in DMEM for 30 minutes at 37°C and then inoculated with Neth/09-*Renilla* virus at an MOI of 3 for 1 hour at 37°C in the presence of the inhibitors. Thereafter, cells were maintained in post-infection DMEM supplemented with NH_4_Cl (25mM) and 6µM Renilla luciferase substrate. Luminescence measurements were acquired at the indicated times post-infection with a Perkin Elmer plate reader. Mock-infected cells were utilized to determine background luminescence.

### Cell viability

The indicated cell lines were treated with inhibitors (Dynasore or Pitstop 2) or controls (NH_4_Cl or DMSO) as described above. Thereafter, cells were washed once with PBS and maintained in post-infection DMEM supplemented with NH_4_Cl (25mM). Cell viability was determined at 24 hours post-inhibitor treatment with the CellTiter-Glo Luminescent Cell Viability Assay (Promega) as per manufacturer’s instructions. Luminescence measurements were acquired with a Perkin Elmer plate reader.

## RESULTS

### Construction and expression of an antibody-based IAV receptor

We combined anti-HA antibody heavy and light chain binding domains from the 5J8 MAb and the cytoplasmic tail, transmembrane and stalk sequences of fTfR to generate the artificial IAV receptor, fTfR-5J8 (Figs. 1A-E).

To test the expression and functionality of the chimeric receptor in the absence of Sia, we expressed the protein in an *Slc35A1* gene knockout background in two human cell lines: the human embryonic kidney 293 (HEK) cells and the adenocarcinoma human alveolar-derived epithelial cells (A549). Both cell lines are permissive to IAV entry and productive infection (43, 44). Sia expression levels in *Slc35A1* knock out (KO) HEK or A549 cells were confirmed by staining with SNA or MAL I/II lectins via immunocytochemistry (Fig. 2A, D) and flow cytometry (Fig. 2B, C, E, F). SNA recognizes α2,6-linked Sia residues whereas, MAL I/II binds to α2,3-linked Sias. HEK and A549 WT cells, as well the *Slc35A1* reconstituted cells displayed higher levels of either α2,6- or α2,3-linked Sias compared to their *Slc35A1* KO versions. The KO cells exhibited a 95-60-fold decrease in α2,6- and α2,3-Sia levels (Fig. 2B, C, E, F)

**Figure 2.**
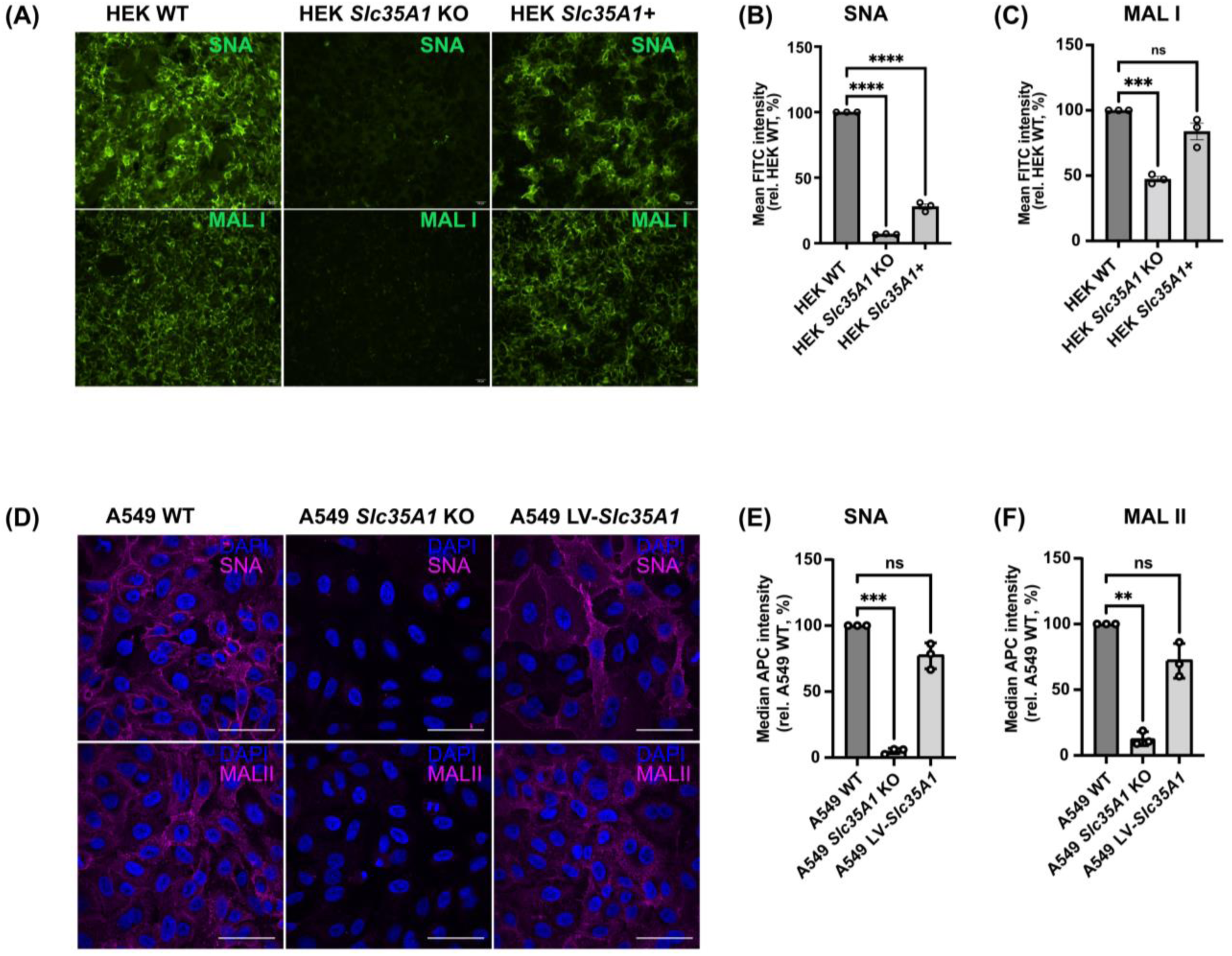
Validating Sia expression in HEK293 and A549 WT and *Slc35A1* KO conditions. (A) Representative fluorescent microscopy images of fixed cells stained with fluorescein-labeled SNA or MAL I. Relative fluorescence level of (B) SNA- and (C) MAL I, stained live cells quantified by flow cytometry. MFI intensities were normalized relative to HEK WT Sia expression (B, C). Background (unstained) subtracted MFI values were analyzed using PRISM software; MFI = Mean fluorescent intensity. *, Error bars show mean ± standard mean error. Statistics were calculated using ANOVA. P<0.05; ****P<0.0001; n=6. (D) Unmodified (WT), and *Slc35A1* KO A549 cells transduced with an *Slc35A1* encoding lentivirus (LV-*Slc35A1*) were fixed and stained with biotinylated SNA or MAL II (magenta) and DAPI (blue). The Alexa Fluor 647-Streptavidin signal intensities were assessed via microscopy. Images are representative of n=3 independent replicates. Scale bar represents 50µm. (E, F) Cells from (D) were incubated with biotinylated SNA or MAL II and the APC-streptavidin in live cells analyzed via flow cytometry. Following subtraction of the median APC signal of corresponding background samples, the median APC signal was normalized to that of WT cells. Data are means +/- standard error of mean/standard deviation from n=3 independent experiments. Statistical significance was inferred by two-tailed one sample t test with a theoretical mean of 100. ***P<0.001, **P< 0.01.

### HA-hFc probe and IAV bind to fTfR-5J8 expressing cells

Surface expression and HA recognition of the fTfR-5J8 receptor were tested by transiently expressing the fTfR-5J8 in HEK *Slc35A1* KO cells for 24hrs, then incubating with an HA-hFc probe comprised of Ca’09 HA fused to a human IgG1 Fc domain. Flow cytometry analysis of the stained non-permeabilized cells showed ∼100-fold increased fluorescence intensity compared to control cells expressing the full length fTfR (Fig. 3A, B). The HEK WT cells also bound the HA-hFc probe, albeit to a much lower level (Fig. 3B). Next, we evaluated the binding of A/Netherland/602/09 virions (99% HA sequence identity to Ca’09) to A549 *Slc35A1* KO cells stably expressing either fTfR-5J8, SLC35A1, or full length fTfR (Fig. 3C). Virions showed significantly higher binding to cells expressing fTfR-5J8 compared to those expressing the full length fTfR control (Fig. 3D, E), thus confirming the expression of the fTfR-5J8 receptor on the cell surface and its recognition of the pandemic H1N1 HA antigen. A549 *Slc35A1 KO* cells stably expressing also showed significantly greater virus binding, as expected (Fig. 3D-E).

**Figure 3.**
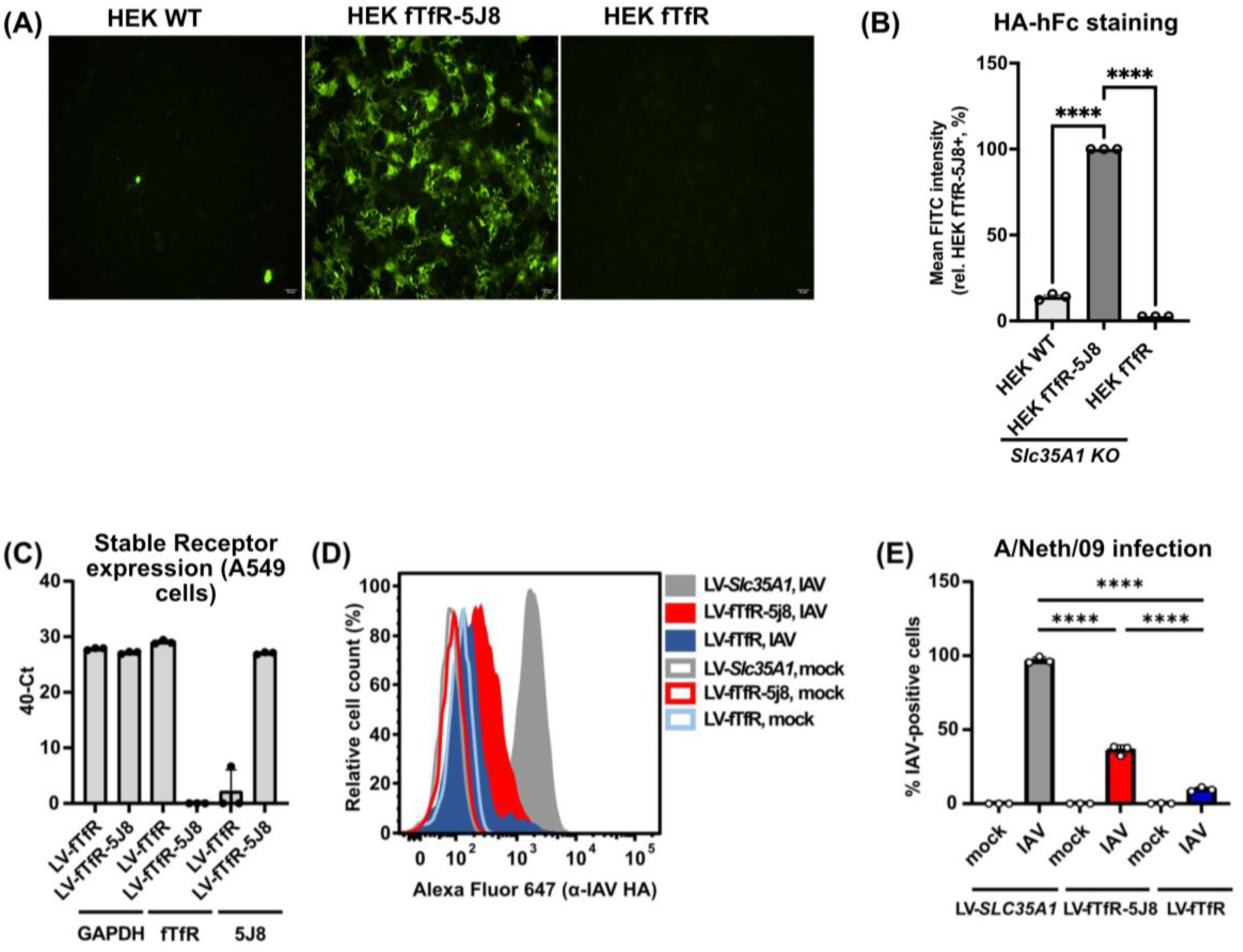
Binding analysis of HA Probe or live virus to Antibody-based receptor expressing cells. (A) Representative fluorescent microscopic images of fixed cells incubated with HA-hFc probe and detected with fluorescein-labelled anti-human Fc. (B) Relative fluorescent intensities of live HA-hFc-stained cells quantified by flow cytometry. Fluorescent intensities were normalized relative to fTfR-5J8 expressing cells. (C) The fTfR and fTfR-5J8 expression was analyzed in A549 *Slc35A1 KO* cells transduced with lentiviruses encoding fTfR (LV-fTfR) or fTfR-5J8 (LV-fTfR-5J8) via qPCR. Primers targeting GAPDH, the extracellular domain of fTfR and 5J8 were utilized. Data are 40-Ct values from technical triplicates from n=1 experiment. A Ct of 40 was assigned to samples where no amplification was detected. (D) Cells from (C) and A549 *Slc35A1* KO cells stably expressing SLC35A1 (LV-*Slc35A1*) were inoculated with A/Netherlands/602/09 at a MOI 25 for 1.5hr on ice. Cells were incubated with an anti-HA antibody and the signal from bound virus quantified via flow cytometry. A representative histogram from n=3 independent experiments is shown. (E) Quantification of the percentage of IAV positive cells from (D). The gating strategy was established using the mock-infected sample. Data are means ± standard deviation and statistical significance was inferred by one-way ANOVA with Sidak’s multiple comparisons test. ****P<0.0001.

### The fTfR-5J8 receptor rescues viral infection in Sia deficient cells

To test whether IAV can infect cells after binding to the fTfR-5J8 receptor, we inoculated the fTfR-5J8 expressing or control HEK cells with virus at a multiplicity of 0.2 and incubated the cells for 8hrs (Fig. 4A), which corresponds to one full replication cycle (45). Results are shown as fluorescent antibody staining for the NP protein (Fig. 4A, B), and percentage of cells infected, which was normalized to the percentage seen for the WT control cells. The fTfR-5J8 receptor rescued to within ∼70% of Ca’09 WT infection in the *Slc35A1* KO HEK cells (Fig. 4B). Cells transiently reconstituted by expression of *Slc35A1* from a plasmid displayed similar levels of IAV infection to HEK WT cells, while no Ca’09 infection was detected in the control fTfR expressing cells (Fig. 4B). To assess the specificity of the fTfR-5J8 receptor, we asked whether it could rescue H1N1s other than those of pandemic origin to similar levels as the pandemic H1N1s tested. We tested this using the laboratory adapted H1N1 strain A/PR/8/1934 (PR8), and found that the fTfR-5J8 receptor did not rescue infection of PR8 HEK *Slc35A1* KO cells (Fig. 4C).

**Figure 4.**
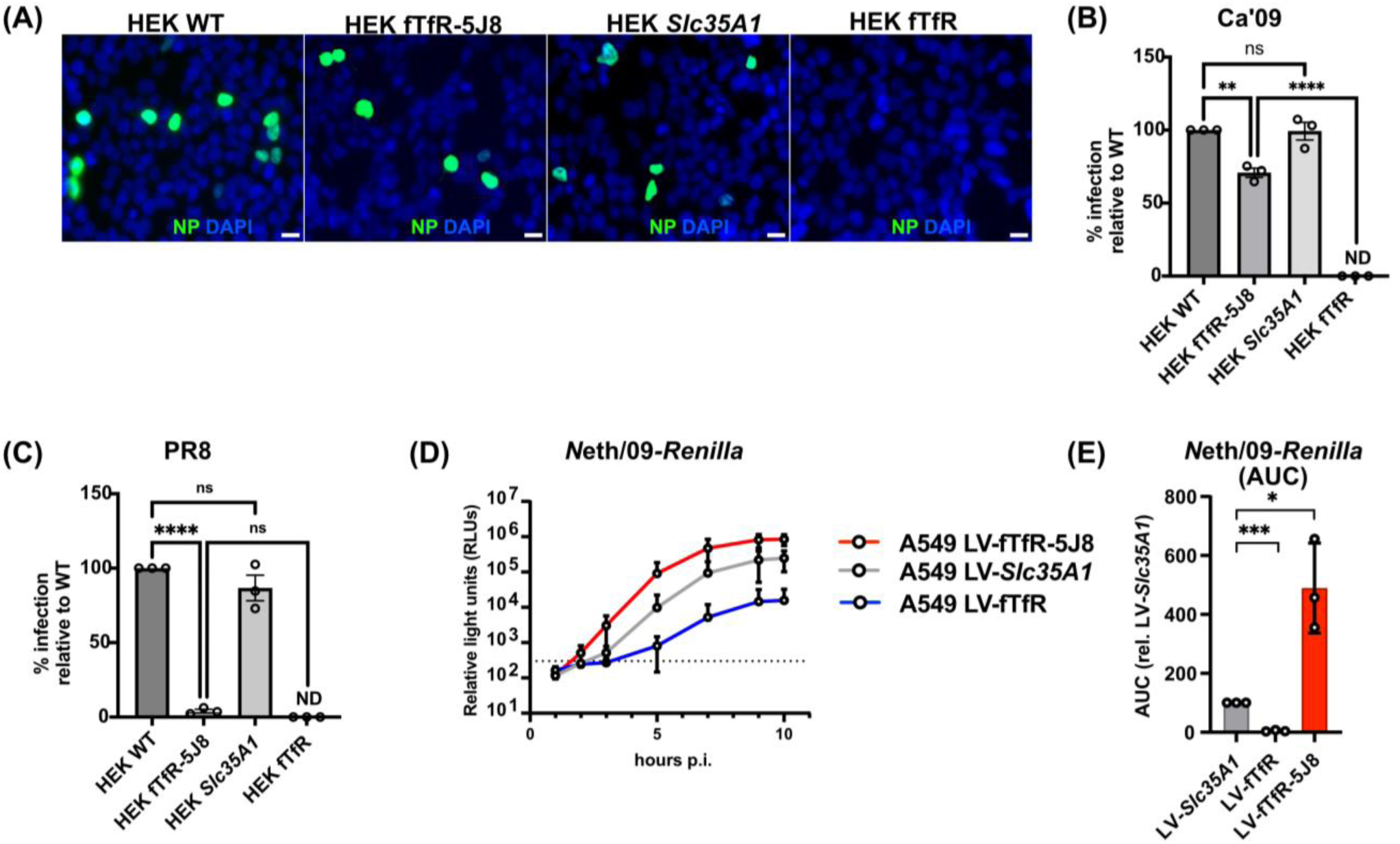
Antibody-based receptor rescue IAV infection in the absence of Sia. (A) Representative fluorescence microscopy images of fixed Ca’09 infected cells. Receptors (fTfR-5J8 and full-length fTfR) were transiently expressed on HEK *Slc35A1 KO* for 24hrs before viral infection (MOI = 0.2) for 8hrs. Cells were fixed and stained with anti-NP antibody (green) and DAPI (blue). (B and C) Infection data for Ca’09 or PR8 infected cells respectively. Four different fields of view were imaged for each condition and % infection was determined. Data was normalized to HEK WT infected cells. (D) Infection curve for the Neth/09*-Renilla* at MOI = 3 in A549 LV-fTfR, LV-fTfR-5J8 and LV-*Slc35A1* cells. (E) AUC plot from D, where AUC values of LV-fTfR-5J8 and LV-fTfR are shown relative to LV-*Slc35A1.* ND = No infection detected, NS = not significant, AUC = Area under the curve, MOI = Multiplicity of infection. All experiments were performed in three independent replicates (n=3). Error bars show mean ± standard mean error. Statistics were calculated using ANOVA. *P<0.05; **P<0.01, ****P<0.0001.

We further examined functionality of the artificial receptor by infecting the *Slc35A1* KO A549 cells transduced to express fTfR-5J8, fTfR or SLC35A1 with the pandemic H1N1 A/Netherlands/602/2009 strain expressing the *Renilla* luciferase gene (Neth/09-*Renilla*) as previously described in (33). Live cell luminescence readout of the various A549 cells inoculated with Neth/09-*Renilla* over the course of 10hrs at MOI 3 showed results similar to that of the Ca’09 infection (Fig. 4D). Relative luciferase unit (RLU) and AUC plots of the infected fTfR-5J8 expressing cells were ∼80-90-fold greater than those of the control fTfR expressing cells, and similar to *Slc35A1* reconstituted LV transduced A549 cells (Fig. 4D, E).

### Antibody-based receptor uptake employs similar endocytosis pathway as Sia-mediated entry

As a major IAV entry route in the absence of serum is the clathrin-mediated endocytosis (CME), we next investigated whether this was the case in the antibody-based receptor system. A549 *Slc35A1* KO LV-*Slc35A1* and LV-fTfR-5J8 cells were treated with inhibitors targeting dynamin (Dynasore) or CME (Pitstop 2) and then infected with A/Neth/09-*Renilla* virus in their presence. Inhibitor treatment did not impact cell viability (Fig. 5A) and led to a dose-dependent decrease in viral replication in cells with reconstituted sialic acid expression and those overexpressing the antibody-based receptor (Fig. 5B-C). In summary, IAV entry mediated by the fTfR-5J8 receptor is dependent on dynamin and CME, akin to that in a Sia-positive context.

**Figure 5.**
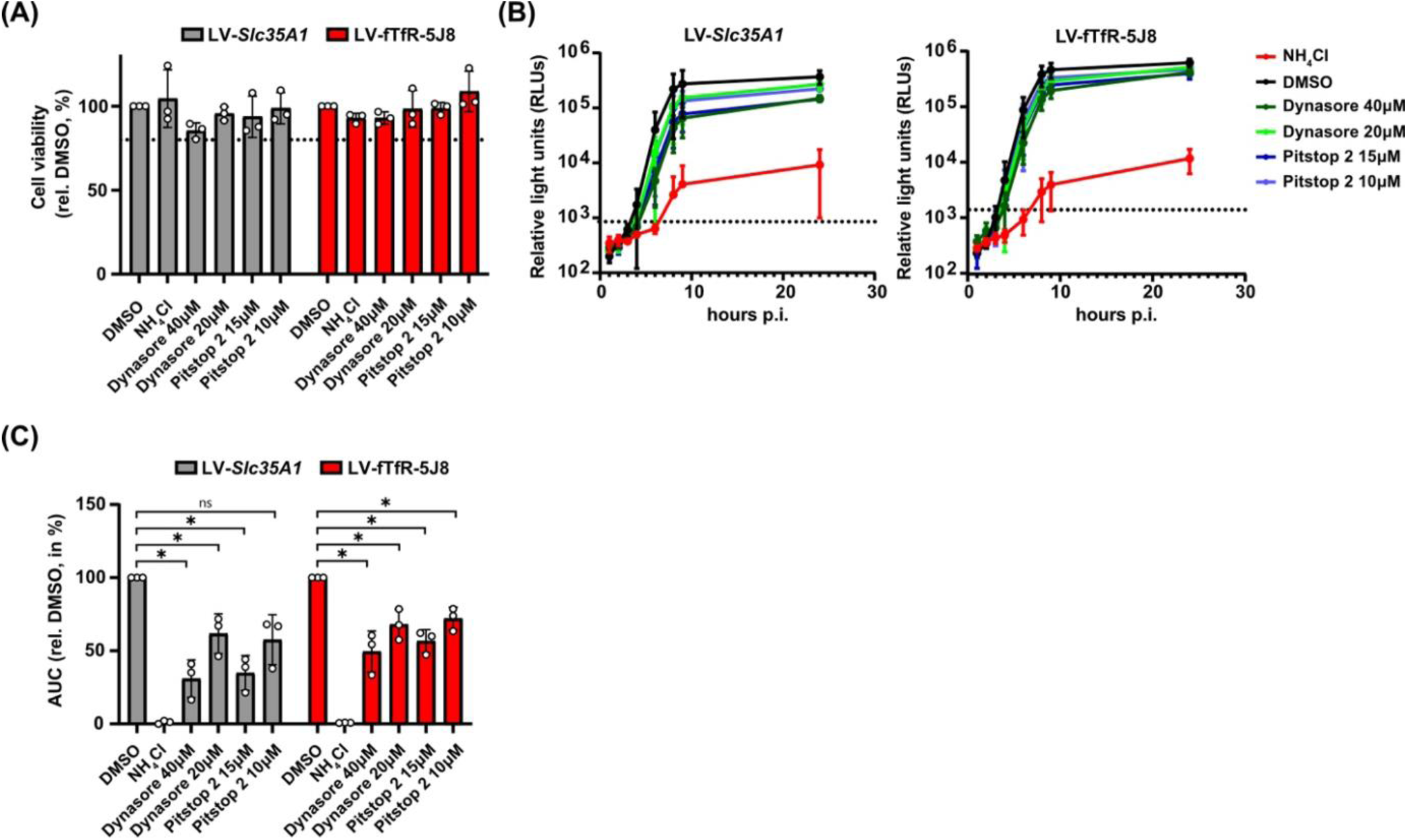
IAV entry mediated by the antibody-based receptor is dynamin and clathrin-dependent. A549 *Slc35A1* KO cells stably expressing SLC35A1 (LV-*Slc35A1*) or fTfR-5J8(LV-5J8-TfR) were pre-treated with Dynasore (40-20µM), Pitstop 2 (15-10µM), NH_4_Cl (25mM) or DMSO (0.15%) for 30 minutes at 37°C and then infected with Neth/09-*Renilla* at a MOI 3 in their presence. (A) Cell viability at 24 hours post-inhibitor treatment relative to the DMSO control. (B) Neth/09*-Renilla* infection curves. Dashed line indicates luminescence from the mock-infected sample. (C) AUC values from (B) were normalized to the DMSO-treated sample within each cell line. (A-C) Data are means ± standard deviation from n=3 independent experiments. Statistical significance in (C) was determined by two-tailed one sample t-test with a theoretical mean of 100. *P<0.05, ns = not significant.

**Figure 6.**
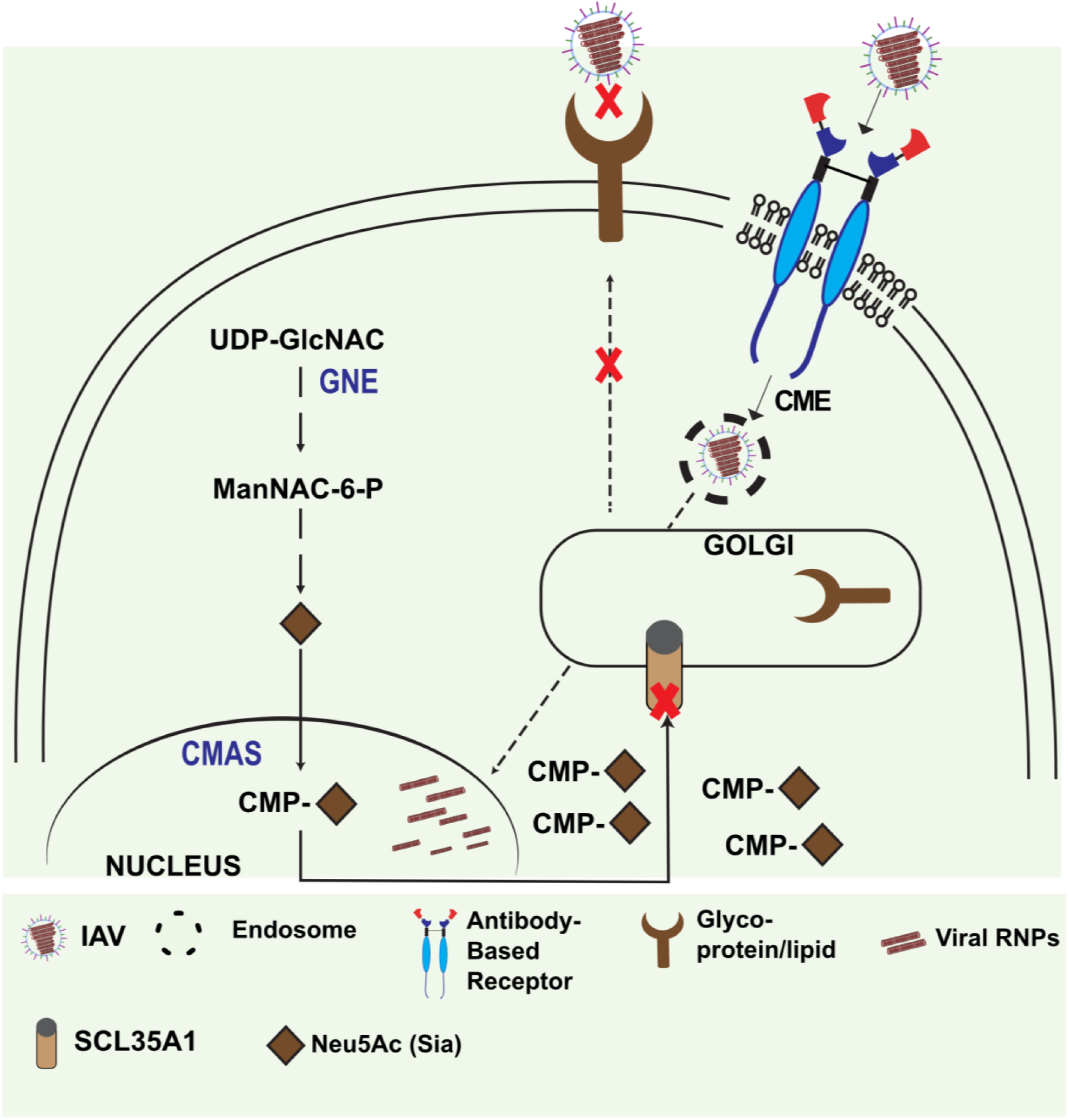
Summary of the new receptor prepared here, and its potential for analysis of the IAV entry pathways. HEK and A549 *Slc35A1* KO cells were prepared, allowing us to examine the rescue of IAV uptake, internalization, and infection in the absence of efficient Sia surface expression by using the alternative receptor. Schematics of the Sia biosynthetic pathway is shown (summarized in (60) indicating the accumulation of CMP-Sia in the cytosol due to the gene deletion of the antiporter SLC35A1. CME, Clathrin mediated endocytosis; UDP-GlcNAC, uridine diphosphate N-acetylglucosamine; ManNAC-6-P, N-acetyl-mannosamine 6-phosphate; Neu5AC’ N-Acetylneuraminic acid; CMP, cytidine monophosphate; GNE = UDP-GlcNAc 2-epimerase/ManNAc-6-kinase.

## DISCUSSION

Here we have developed a new tool for the study of influenza virus binding, entry and infection of cells, by preparing a glycoprotein receptor that binds specifically to the same region of HA as Sia receptors. By using a protein ligand that combines an antibody as a defined binding structure and a truncated receptor with a well-understood cell entry pathway involving clathrin-mediated endocytosis, we can use the fTfR-5J8 receptor to both understand and manipulate the processes involved. For Multiple IAVs, studies over the past 70 years have shown that viral infection requires Sia binding (46–49). It is also clear that differences in the affinity of Sia binding are directly correlated with the efficiency of infection, for example those seen between the α2,3- and α2,6-Sia linkages found in birds and humans respectively (50, 51). However, we lack a clear understanding of how that binding leads to uptake or entry and infection (52). It is difficult to modify Sia receptors on cells or in animals, as the Sia, oligosaccharides, and glycoproteins or glycolipids that form the functional receptors are the products of diverse and variable series of enzymatic activities, making it difficult to change the receptor properties with any precision or predictability.

The fTfR-5J8 receptor combines the HA-binding domains as an scFv of an anti-pandemic H1N1 antibody (MAb 5J8) fused to the cytoplasmic tail, transmembrane and stalk domain of the fTfR. Uptake of the TfR and its regular ligand holo-(iron-bound) Tf occurs almost entirely via CME due to the efficient engagement of the TfR cytoplasmic tail with the adaptor protein 2 (AP2), which links it to clathrin-coated pits (53, 54).

The fTfR-5J8 was expressed on the cell surface where it bound to the viral HA as an HA-hFc fusion protein, and it also efficiently rescued infection on both HEK293 and A549 cells that lack Sia cell surface expression due to knock out of the Sia-CMP transporter SLC35A1 (55). The fTfR-5J8 expression cells also showed comparable levels of IAV infection rescue to those seen in SLC35A1 reconstituted cells. The fTfR-5J8 receptor bound to the virus appears to follow a similar uptake pathway as has been seen in Sia-expressing cells, as evidenced by the comparable sensitivity to dynasore and pitstop, inhibitors of dynamin and clathrin-mediated endocytosis, respectively.

Future studies will allow us to closely examine the entry and infection pathways of the H1N1 pandemic viruses by manipulation of the properties of the fTfR-5J8 receptor. For example, mutation of the scFv-binding site (or of the HA epitope) would reveal the effects of lower (or potentially higher) affinity interactions. Mutagenesis of key sequences within the TfR cytoplasmic tail will allow the the uptake and entry pathways to be manipulated, for example so that the receptor may traffic into alternative endosomal pathways (35, 56).

Other aspects of the viral infection process might also be revealed, including a better understanding of the role of the NA and cell surface binding in the viral cell infection and release pathways (57–59). While some functions of NA in viral budding will not be required - HA-Sia crosslinking of newly created particles will not occur during budding - the presence of the fTfR-5J8 on the cell surface may bind to the virus or to free HA trimers either in the endoplasmic reticulum, Golgi, or at the cell surface. The fTfR-5J8 likely has a higher affinity for HA than Sia, but we can test the effects of different levels of receptor expression in cells by mutagenizing the scFv binding structures to alter their affinity or avidity for the HA. Thus, this receptor serves as a versatile system for studying IAV uptake and internalization mechanisms in a controlled manner.

## ACKNOWLEGEMENTS.

Kirsten Young and Brynn Lawrence assisted in the development of the system, and Patrick Wilson, Weill Cornell University provided the antibody constructs used to prepare the scFv used here. We also thank Elisabeth Gaggioli and Alena Iseli for technical assistance.

